# Diversity patterns and selective sweeps in a Southeast European panel of maize inbred lines as combined with two West European panels

**DOI:** 10.1101/2020.11.24.376087

**Authors:** Vlatko Galić, Violeta Anđelković, Natalija Kravić, Nikola Grčić, Tatjana Ledenčan, Antun Jambrović, Zvonimir Zdunić, Stéphane Nicolas, Alain Charcosset, Zlatko Šatović, Domagoj Šimić

## Abstract

More than one third of European grain maize is produced in South Eastearn Europe (SEE) and utilization of historical maize material developed in SEE for its favorable alleles and diversity has long been speculated. However, molecular diversity of the SEE maize genetic material is scarce. The objectives of this study were i) to analyze diversity patterns in a large panel of densely genotyped historical accessions from SEE, ii) to compare the data with those obtained from other two European panels, and iii) to identify genomic regions that have undergone selection (selective sweeps) in response to adaptation to SEE conditions. 572 accessions of the historical inbred lines from Maize Research Institute Zemun Polje representing the SEE material were genotyped using the 600k maize genotyping Axiom array. The genotyping results were merged with two European panels DROPS and TUM. Genetic structure and diversity were analyzed using neighbor-joining cladogram, PcoA, Admixture, Structure and sNMF. To detect the selective sweep signals, Tajima’s *D* statistic and RAiSD were employed. The best number of ancestral populations was K=7, whereby one of them is a subpopulation containing inbreds belong exclusively to the SEE panel. The prevalence of inbreds linked to historical US inbred lines Wf9, Oh43, Pa91 and A374 was detected in SEE. Possible soft selective sweep was detected in chromosome 2 in region harboring a gene linked to promotion of flowering FPF1. Additional scan for selective sweeps using the RAiSD methodology yielded four signals in chromosomes 5 and 6, all in gene-rich regions. Several candidates of selection were identified, influencing the plant morphology and adaptation. Our study provides the first step towards the utilization of the SEE genetic materials for use in maize breeding. Phenotypic analysis is needed for assessment of SEE accessions for favorable alleles, and identification of breeding targets.

## 1. Introduction

Maize (*Zea mays* L.) breeding is based on the selection of the favorable progenies from the designed crosses between inbreds bearing favorable alleles/favorable genetic background (Hallauer et al., 2010). This type of advanced-cycle pedigree breeding scheme might lead to the available maize germplasm becoming more elite, although more genetically narrow (Lu and Bernardo, 2001; Reif et al., 2005). Due to the distinct heterotic patterns in maize breeding (Lee and Tracy, 2009), population-level diversity is maintained, but to sustain the long-term breeding progress, exploiting of the new germplasm resources is inevitable, especially for adaptation traits (Bouchet et al., 2013; Romero Navarro et al., 2017; Wegary et al., 2019).

Modern maize hybrids grown around the world today, are mostly single crosses developed through tangled crossing and testing schemes in target populations of environments by multi-national companies. Only marginal market shares are held by the small companies and public institutions. The global seed market can be separated into two tiers. The first tier represents 10 largest companies owning 69% of the world market and only three of which reported sales of >3000 $m for 2018, and the other represents all other stakeholders. This trend can be easily extrapolated to maize only, especially due to the fact that maize seed business accounts for 42% of the global seed market of all crops with global sales of nearly 18 billion US$ in 2018 (FAO/IHS Markit Agribusiness Consulting, 2019).

However, the evolution of the seed business was driven by the evolution of maize breeding itself, initially mainly through the public breeding programs. During the 19^th^ century, the US corn market was prevailed with seeds of many open pollinated varieties (OPVs) adapted to temperate environments from several early breeding programs, such as Reid Yellow Dent, Lancaster Sure Crops, Leaming corn, etc. By the year 1933, first significant acreages of the double cross hybrid corn were reported (Troyer, 2009) with substantially higher yields compared to OPVs. Further developments in hybrid breeding were observed, especially with development of Stiff Stalk Synthetic during the 1930s. From the 1950s there was a rapid shift from breeding corn by farmers for farmers, to breeding by seed companies which led to further increase in grain yields. Interestingly, 87% of the maize genetic material utilized in U.S. during the mid-2000s could be traced back to only five historical OPVs, with highest leverage of the variety adaptness to surpass thousands of other, today-probably extinct OPVs (Arca et al., 2020; Troyer, 2004).

European perspective on the maize breeding was somewhat different than the US one. Some of the first introductions of maize into the parts of Europe after the discovery of Americas probably failed on the wider scale, due to the low levels of adaptation to European climatological conditions. It is well established that during the early 16^th^ century, several populations of Caribbean origin were widespread in southern Spain and Italy, but it was probably not until the separate introductions of the Northern Flints later in the same century, that the maize has been broadly adapted to European mid-latitudes (Mir et al., 2013; Rebourg et al., 2003; Tenaillon and Charcosset, 2011).

South Eastern Europe (SEE), consisting primarily of the Balkan Peninsula, can be considered as a European counterpart to the US Corn Belt with well adapted late temperate germplasm and more than 20% of the crop areas under maize (Leff et al., 2004). Moreover, more than 35% of the European grain maize was produced in Serbia, Romania and Hungary and continental Croatia in the period from 2010 – 2014 (USDA, 2020). In more recent reports, Croatia, Serbia, Romania and Hungary in 2018 and 2019 together contributed 52% and 51%, respectively of the European Union + Serbia total maize grain production (Eurostat, 2019; Republic of Serbia, 2020).

In the former Yugoslavia, a large number of the local varieties (>2000) classified into 18 races, showed large within-race and among-race variability and expected heterozygosity (Geric et al., 1989; Ignjatović-Micić et al., 2013) probably reflecting the multiple origins and introductions of maize to these areas also seen in words in different languages designating maize as Turkish corn, or “kolombač” a word straining from the word Columbus in Montenegro (Leng et al., 1962). Based on the morphological assessment, landraces of the former Yugoslavia resemble many different historical populations such as Amarillo de Ocho (Small-ear Montenegrin flints), US Northern Flints (Eight-rowed flints), Old Southern Dents (Many-rowed Soft Dents), etc. along with several more recent OPVs from late 19^th^ century such as Hichory King (Large kernel dents), and early 20^th^ century introductions of Golden Mine and Queen of Prairie (Rumski zlatni zuban) (Andjelkovic et al., 2012; Babic et al., 2012; Kozumplik and Martinić-Jerčić, 2000).

After the World War II, some of the European traditional varieties were used to develop hybrids adapted to European conditions (Tenaillon and Charcosset, 2011), and were crossed to materials developed from the US imported double-cross hybrids such as WF9 x Hy, Hy x Oh07, W32 x W187, etc. during the 1950s (Brkić et al., 2003; Hadi et al., 2013). Growing the locally bred maize hybrids was so popular in the SEE during the 1960s, that it was even speculated to surpass the production of the US hybrids in the following decades (Leng et al., 1962). The source of that-time modern introduced US germplasm was the organized production of US double cross hybrids in Yugoslavian public research institutes as part of the American Aid plan through the Foreign Organization Administration from the original inbreds (Tavčar, 1955). The imported inbreds were: Wf9, 38-11, Hy, L317, N6, K148, K150, M14, W32, W187, A374, A375, and Oh07.

Data about molecular diversity of maize genetic material in SEE is scarce (e.g. Şuteu et al. 2013 (Şuteu et al., 2013)). Nonetheless, utilization of the SEE maize for its favorable alleles and diversity has been long speculated (Leng et al., 1962), with most of the materials still deposited in gene banks. One such bank is Maize Research Institute Zemun Polje (MRIZP) gene bank conserving >6000 accessions, of which >2000 are the maintained local landraces collected throughout the former Yugoslavia and > 2000 accessions are the inbred lines originating from 40 different countries (Vančetović et al., 2010) representing one of the largest maize collections in the world (Gouesnard et al., 2017). The view on the relevance of the plant genetic resources has at least two converging aspects. First is the conservation of the biodiversity that has been narrowed by the way the historical diversity has been utilized (Planchenault and Mounolou, 2011). The other aspect is to use all available modern breeding tools such as dense genotyping, high throughput phenotyping, etc. to mine and utilize the favorable variability by overcoming the issues such as linkage drag (Hölker et al., 2019; Ortiz et al., 2010; Sood et al., 2014; Unterseer et al., 2016).

The objectives of this study were i) to analyze diversity patterns in a large panel of densely genotyped historical accessions from SEE, ii) to compare this genetic diversity with two European diversity inbred line panels, and iii) to identify genomic regions that have undergone selection (selective sweeps) in response to adaptation to SEE conditions.

## 2. Material and Methods

### Plant material

The 572 accessions of the Maize Gene Bank of the Maize Research Institute Zemun Polje (MRIZP) were used to carry out this study. Accessions i.e. inbred lines were chosen in a way to represent the diversity of introduced or de-novo developed material from the SEE breeding programs along with several inbreds with collection attributes from other countries. In the SEE panel, there were 220 accessions collected from Bulgaria, 132 from ex-Yugoslavia, 54 from Romania, 42 from Hungary,18 from ex-Czechoslovakia, 13 from Poland, 7 from Greece, along with inbreds that did not originate from SEE: 47 from ex-USSR, 12 from USA, 8 from Mexico, 7 from Iran, 3 from France, 2 from Canada, 2 from ex-East Germany, 1 from ex-People’s Republic of Korea, 1 from Pakistan, 1 from Switzerland, 1 from Argentina and 1 of unknown origin. All additional information about the used inbred lines is available as Supplementary table S1.

### Genotyping and data management

The MRIZP accessions of the SEE panel were genotyped with Axiom™ 600k Maize SNP Genotyping Array with 616,201 variants of which 6,759 represent insertions/deletions (Unterseer et al., 2016, 2014). All steps of the DNA analysis were conducted by TraitGenetics GmbH, Germany including standard protocols of DNA extraction and marker quality control. Two other publically available genotypic matrices anchored with the same genotyping array were used to conduct this study. First was data from Unterseer et al. (Unterseer et al., 2016) on 155 elite Dent or European flint / Northern Flint inbred lines, mainly from German and French public breeding programs (TUM panel) and the second was the data from Millet et al. (Millet et al., 2016) on 247 dent inbred lines (DROPS panel). Most of the inbred lines from both data sets were European developments. Additional information about the inbred lines is available in Supplementary table S1.

The data from all three datasets were merged using a custom R script and insertions/deletions were removed, leaving 500,167 overlapping positions. Positions were further filtered for heterozygotes (2.5%) and missing data (5%) in Tassel software (Bradbury et al., 2007) version 5.2.64 leaving a final set of 460,243 filtered positions. The positions were imputed using the LinkImpute method (Money et al., 2015) with 50 sites in high linkage disequilibrium and 30 nearest neighbors. For population structure analysis, all positions were thinned to 1000 base pair distance, leaving 166,755 sites.

### Population structure

Population structure was determined by combining several methods: Neighbor-joining cladogram was constructed in Tassel and edited using a FigTree software (Rambaut, 2018) version 1.4.4.

Principal coordinate analysis (multi-dimensional scaling, PcoA) was performed with thinned marker set with identity-by state distance matrix as input in Tassel software version 5.2.64. To correctly infer the underlying genetic structure of the assessed germplasm, Admixture analysis was run (Alexander and Lange, 2011) in Ubuntu 20.04 terminal with 166,755 imputed and thinned sites. The cross-validation error did not reach minimum until the maximum number of 15 infered populations. Another method for inference of ancestry was sparse nonnegative matrix factorization algorithm (sNMF) (Frichot et al., 2014) in which cross-entropy criterion was employed to find the best value of K, but similarly to Admixture cross-validation results, minimum was not reached until the last assumed ancestral population.

To infer the optimal number of ancestral populations (K), 10,000 positions were randomly sampled from the imputed and thinned set of 166,755 sites and analyzed with STRUCTURE software (Pritchard et al., 2000), version 2.3.4. The K was set from 1 to 15 and 5 runs were carried out per each K with 5,000 burn-in cycles and 15,000 replicates. Based on the findings of Puechmaille (Puechmaille, 2016) that uneven sampling of subpopulations leads to underestimates of true number of K, parameters MedMed K, MedMeanK, MaxMed K and MaxMean K were calculated using the StructureSelector software (Li and Liu, 2018). Additionally, parameter deltaK (Evanno et al., 2005) was calculated.

Spatial projections of the calculated ancestry coefficients were performed using a BioconductoR package LEA (Frichot and François, 2015). Pie charts of the average ancestries of samples with assigned putative origin were mapped to 15 European locations, and Kriging on dominant spatial patterns was performed. Single coordinates were added to each country of origin, while the historical inbreds from ex-East Germany were assigned to Germany pool, ex-Czechoslovakian inbreds were mapped between today Czech Republic and Slovakia and ex-Yugoslavian inbreds were mapped on Serbian-Croatian border harboring the largest ex-Yugoslavian breeding programs.

### Parameters of genetic diversity and selective sweeps

Nucleotide diversity *π* was assessed as 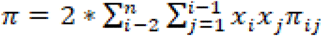 where *x_i_* and *x_j_* are the respective frequencies of the *i*-th and *j*-th sequences, *π* is the number of nucleotide differences per nucleotide site between the the *i*-th and *j*-th sequences, and *n* is the total number of sequences in the sample. Watterson estimator *θ* was calculated as *θ* = 4*N*_*∊*_*μ*, where *N*_*∊*_ is effective population size, and *μ* is an estimate of per-generation mutation rate.

Tajima’s *D* was calculated from the afore mentioned parameters as 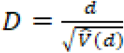, where *d* represents difference between two values of *θ*, and 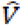 is a variance of this difference. Scan for selective sweeps was carried out using a sliding window analysis with a step size of 100 bp, and a window size of 500 bp in Tassel software version 5.2.64. Another, more stringent protocol for sweep detection was also carried out, namely Raised Accuracy in Sweep Detection (RAiSD) (Alachiotis and Pavlidis, 2018). In RAiSD protocol, three signatures of selective sweeps are calculated: first signature is the local reduction of the polymorphism level quantified by parameter *μ*^*VAR*^. The second signature shows the shift in the site frequency spectrum (SFS) toward low- and high-frequency derived variants and is termed *μ*^*SFS*^. The third signature (*μ*^*LD*^) shows a localized pattern of linkage disequilibrium (LD) levels, characterized by high LD on each side of a putative mutation and low LD between loci that are located on different sides of the beneficial allele. The final parameter *μ* is calculated from the three above mentioned parameters. Window size in the analysis was set to 50 base pairs. RAiSD software version 2.8. (Alachiotis and Pavlidis, 2018) was run in the Ubuntu terminal using the full imputed SNP matrix.

## 3. Results

### Genotyping data summary

In the filtered and imputed dataset, there was a total of 460263 SNPs, 460241 of which were segregating. Average minor allele frequency (MAF) was 0.255, and high values of Tajima’s *D* (4.105) were observed. When SEE panel genotyping data was combined with two other European panels with publicly available data for Affymetrix Axiom 600k chip (Millet et al., 2016; Unterseer et al., 2016), all 460263 loci were found to be segregating, with similar MAF and slightly higher Tajima’s *D* of 4.661 (Table 1)

**Table 1.** Summary of genotypic data for the SEE maize panel as well as publicly available genotypic data for the two West European panels of DROPS (Millet et al., 2016) and TUM (Unterseer et al., 2016).

| Panel | Number of inbreds | Number of sites (all panels) | Segregating sites | Average MAF | $\pi$ per bp | $\theta$ per bp | Tajima's $D$ |
| --- | --- | --- | --- | --- | --- | --- | --- |
| SEE | 572 | 460263 | 460241 | 0.255 | 0.340 | 0.144 | 4.105 |
| DROPS | 247 |  | 460242 | 0.245 | 0.333 | 0.164 | 3.264 |
| TUM | 155 |  | 460239 | 0.264 | 0.359 | 0.178 | 3.351 |
| Total | 974 |  | 460243 | 0.255 | 0.346 | 0.134 | 4.661 |

### Number of ancestral populations (K) and admixture analysis

The used methods Admixture and sNMF failed to reach minimum values of cross-validation error and cross-entropy, respectively up to the maximal inferred number of 15 assumed ancestral populations, although the presence of a “knee” was observed in cross-entropy analysis (not shown). STRUCTURE algorithm was run with a random subset of 10,000 markers and gave two conflicting groups of results depending on the employed methodology (Supplementary figure 1). The ΔK method (Evanno et al., 2005) gave an estimate of five ancestral populations, while LnP(K) method (Pritchard et al., 2000) gave an estimate of seven ancestral populations. The third method employed to support the decision on best number of K proposed by (Puechmaille, 2016) converged in all four estimated parameters (MedMed K, MedMean K, MaxMed K and MaxMean K) on value of K=7 (Figure 1) which was used for further analyses. FST values between the assumed ancestral populations can be seen in Table 2. The first population (K1) represents European flints, present in all three assessed panels, but dominant in the TUM panel. Second population (K2) represents parts of the Stiff Stalk Synthetic-derived germplasm, namely B73, and inbreds from Italy present in the DROPS panel. The third population (K3) is represented by the Mo17-related inbreds, i.e. Lancasters. In fourth population (K4) are the lines derived from Stiff Stalk Synthetic, namely B14 and A632. The fifth population (K5) bears lines derived from Wf9, Oh43 and Pa91. Markedly, in sixth population (K6) in samples with population memberships >0.9 are almost exclusively inbreds from SEE panel, except from a single line from USA, namely A374 (historical Minnesota line) which represents historical US germplasm strained from Reid Yellow Dent. The seventh population (K7) was represented with Iodent pool, focused around Iodent progenitor line PH207. Most interesting was the complete lack of Iodent inbreds from the SEE panel with only two inbreds with ancestral coefficients of 0.706 in K7 from Hungary and ex-Yugoslavia (Supplementary table 1).

**Figure 1:**
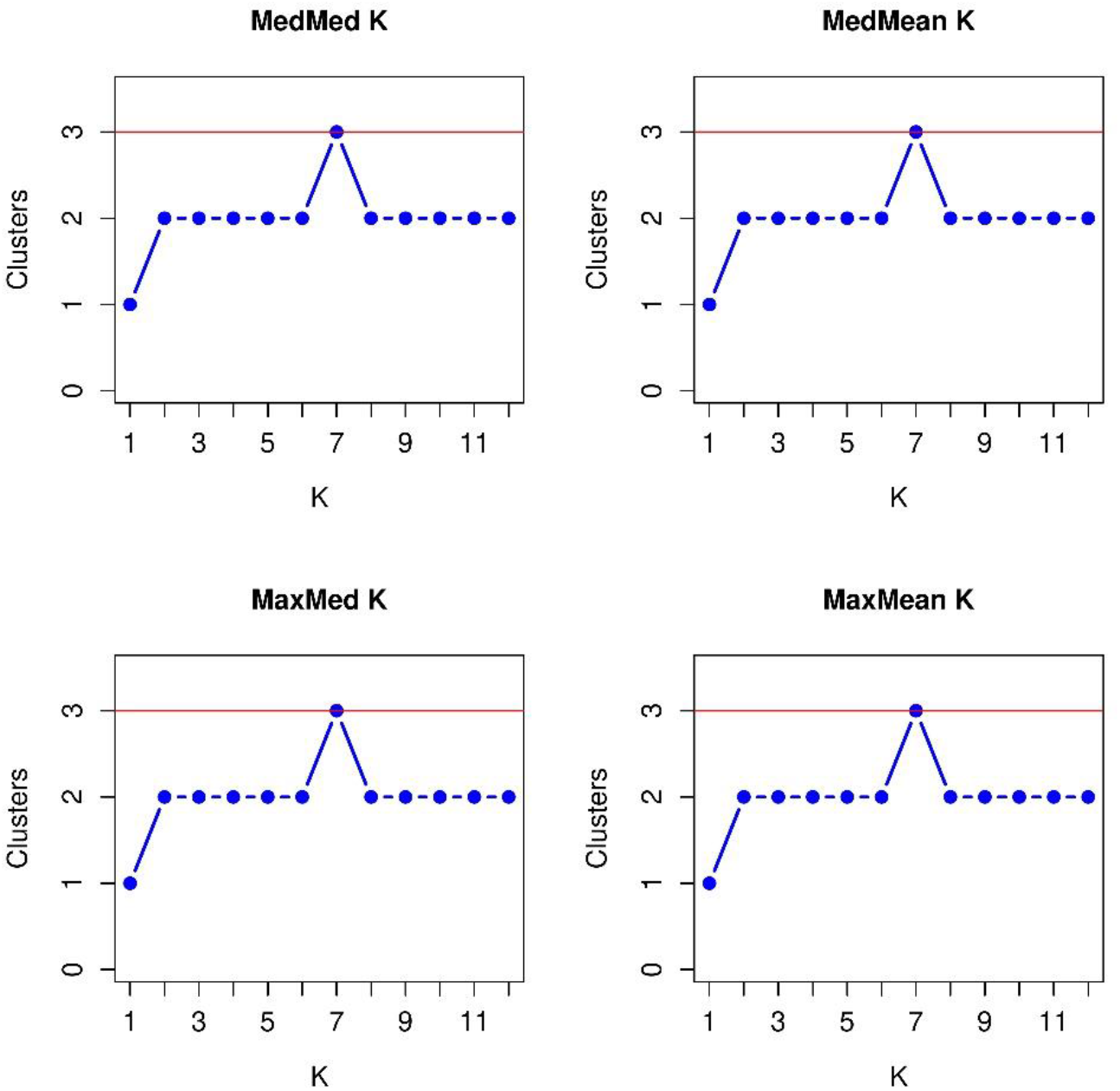
Selection of best number of ancestral populations (red line) using a method developed by (Puechmaille, 2016).

**Table 2.** Mean dissimilarity (FST) between ancestral populations

|  | K1 | K2 | K3 | K4 | K5 | K6 |
| --- | --- | --- | --- | --- | --- | --- |
| **K2** | 0.476 |  |  |  |  |  |
| **K3** | 0.367 | 0.582 |  |  |  |  |
| **K4** | 0.389 | 0.495 | 0.506 |  |  |  |
| **K5** | 0.273 | 0.478 | 0.365 | 0.395 |  |  |
| **K6** | 0.209 | 0.404 | 0.301 | 0.311 | 0.186 |  |
| **K7** | 0.438 | 0.625 | 0.521 | 0.536 | 0.421 | 0.347 |

Based on STRUCTURE results, a highlighted neighbor joining cladogram was constructed (Figure 2). In the cladogram, highlighted are the clades in which inbreds with membership coefficients >0.9 are found. In the cladogram, populations K1, K2, K3, K4 and K7 are distinguished, while individuals of populations K5 and K6 appear scattered on different branches.

**Figure 2.**
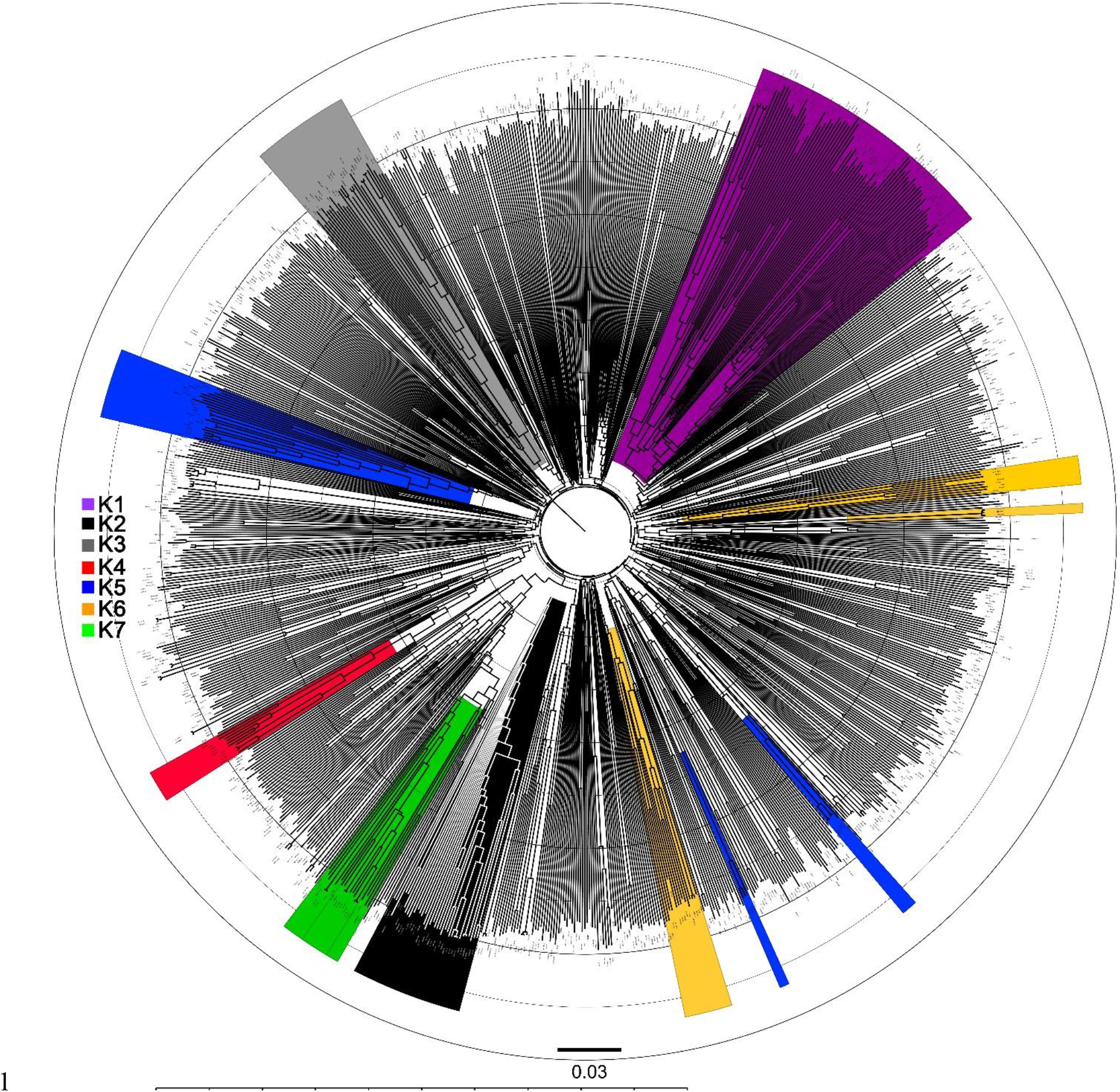
Neighbor joining cladogram of combined three European maize panels SEE, DROPS and TUM (n=974). Highlighted are the clades with inbreds with membership coefficients in admixture analysis >0.9.

Principal coordinate analysis (PcoA)was employed to further analyze the obtained diversity patterns (Figure 3). Compared to the neighbor joining clustering results, only K1 and K7 showed distinct appearance across three examined planes. Lack of distinctness visible between groups with membership coefficients >0.9 K2 and K4, as well as K5 and K6 was partially in agreement with results of neighbor joining clustering. Only the first two coordinates showed eigenvalues >2 (Table 3), with slight decrease in eigenvalues up to the last assumed coordinate (Table 3). Appearance and the spread on the scatterplots (Figure 3) was in accordance with mean pairwise differences in Table 2. Smallest spread was accompanied with lowest observed FST values within populations (Table 2), especially in K2 (0.178) and K7 (0.228)

**Figure 3.**
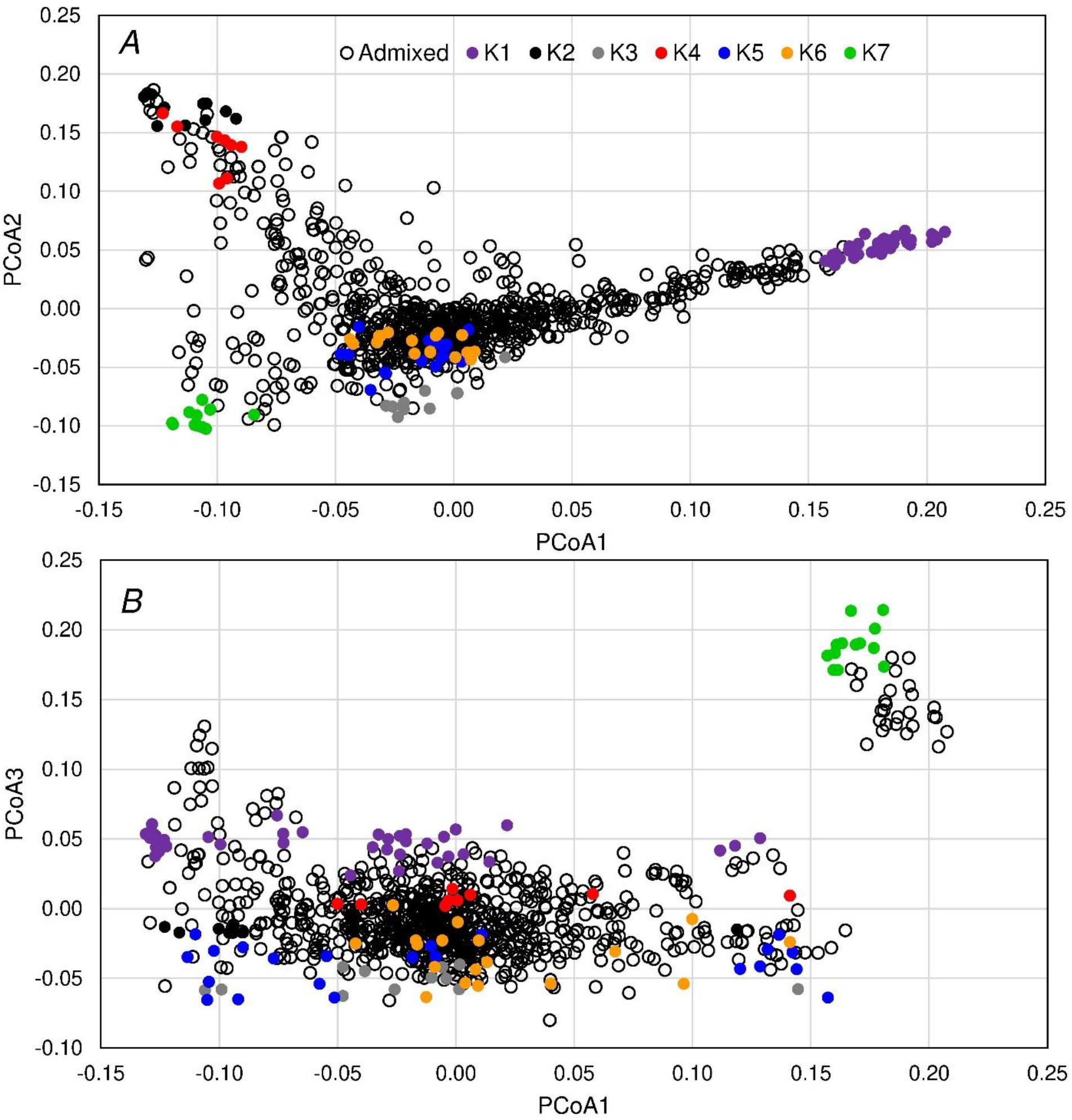
Principal coordinate analysis (PcoA) results of the 974 assessed inbred lines from the three European maize panels. Figure A shows principal coordinates 1 and 2, while principal coordinates 1 and 3 are shown in B. The inbred lines with Admixture membership coefficients >0.9 are shown in color.

**Table 3.** Eigenvalues of the assessed components from the PcoA analysis.

| PcoA | eigenvalue |
| --- | --- |
| 1 | 4.28 |
| 2 | 2.42 |
| 3 | 1.85 |
| 4 | 1.46 |
| 5 | 1.31 |
| 6 | 1.23 |
| 7 | 1.06 |

**Figure 4.**
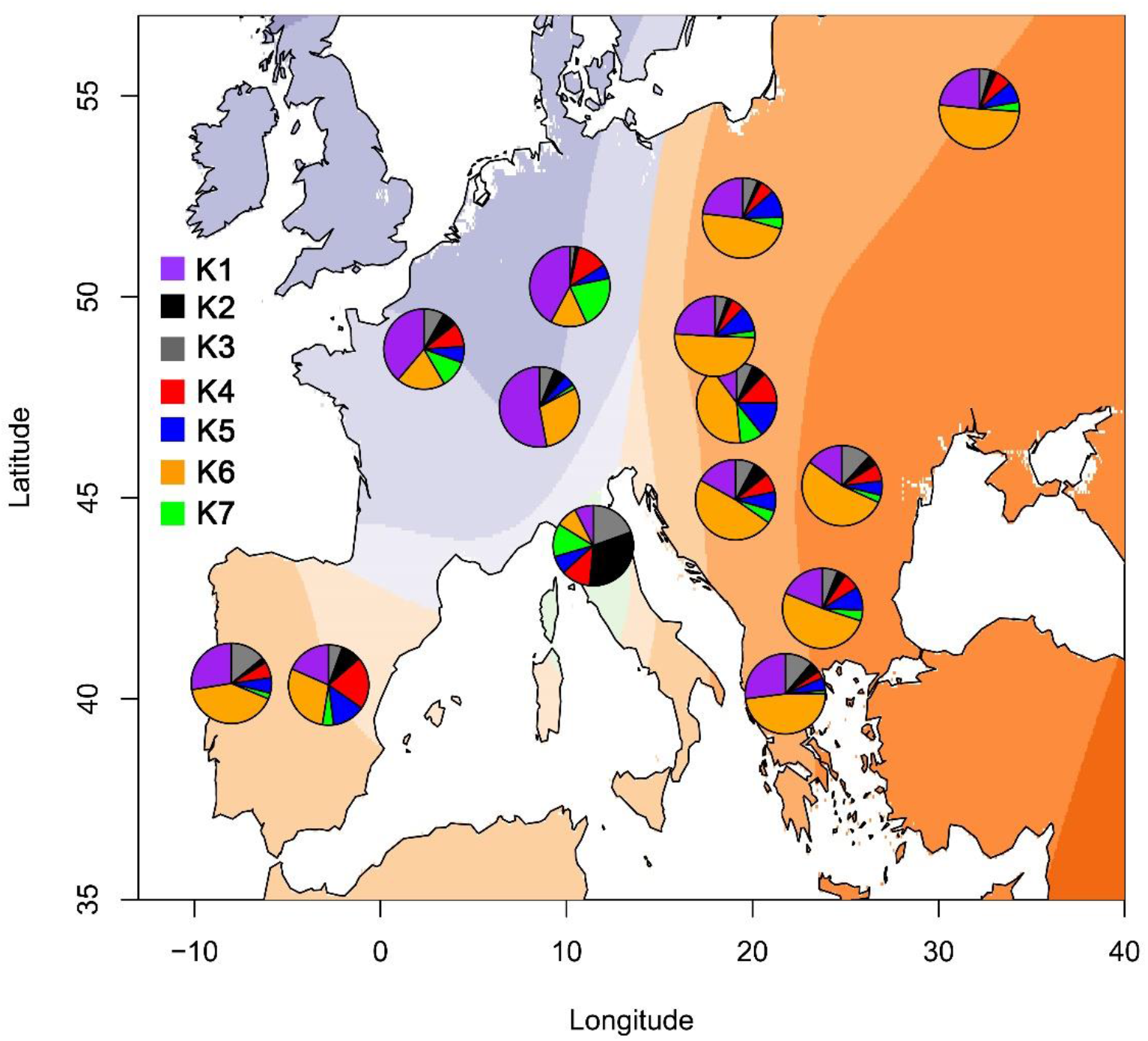
Pie charts of the mean population membership coefficients for the 15 European countries with known or putative origin of inbred lines assessed in the three maize genotyping panels. Different layers of color represent the results of geospatial kriging of the dominant patterns of population membership coefficients.

Kriging of the mean population membership coefficients to 15 known and putative sites of origin of the assessed maize inbred lines showed three different dominant geospatial patterns. First pattern was mostly represented by Germany, France and Switzerland, with prevailing European flint and dent genetic group. The germplasm related to B73 and Mo17 (K2 and K3) was dominantly represented in Italy, while Spain, Portugal, Greece, Bulgaria, Romania, ex-Yugoslavia, Hungary, ex-Czechoslovakia, Poland and ex-USSR showed dominant germplasm from K6 with varying shares of other materials. Higher mean ancestral coefficients linked to historical Minnesota inbreds were also observed in these countries.

### Scans for selective sweeps

In the scan for selective sweeps based on Tajima’s *D* statistics, a single large genomic region with negative values of *D* was detected in the SEE panel on chromosome 2 between 90 and 95 MBp. The negative value of *D* was caused by lower values of parameter π (Figure 5). In this region, on the position 91.2 MBp, a gene coding for Flowering promoting factor-like 1 protein is found. BLAST of the cDNA coding sequence gave 84-100% sequence covers in maize, sorghum, and weeping love grass possibly indicating a conserved gene in C4 grasses.

**Figure 5.**
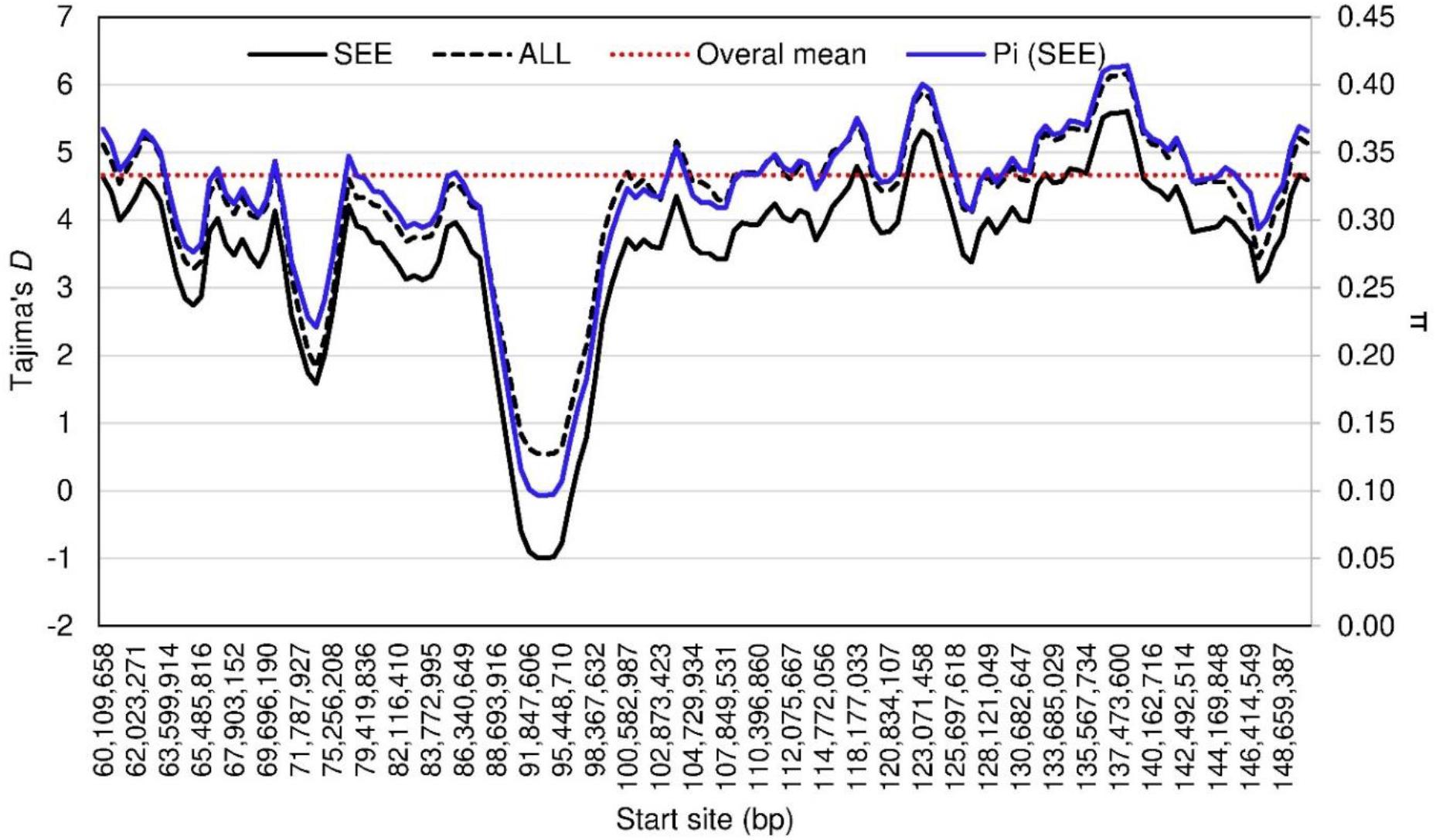
Tajima’s *D* and π values (blue line, secondary axis) for a region on chromosome 2 associated with flowering promoting factor-like 1.

The further scan for selective sweeps was run using the tool RAiSD. The first parameter of the RAiSD analysis μ^VAR^ quantifying the variations per 50 bp window showed most variation around the centromere region of chromosome 5 (Figure 6A). The second parameter μ^SFS^ assessing the shifts from the expected site frequency spectra showed three positions with non-zero estimates (Figure 6B), at start positions 122994833, 144327275 and 164884576, respectively. The μ^LD^ parameter showed expected lower LD values for putative positively selected positions (Figure 6C) resulting in final estimates of sweep statistics (μ) of 0.54, 0.58 and 0.34, respectively (Figure 6D).

**Figure 6.**
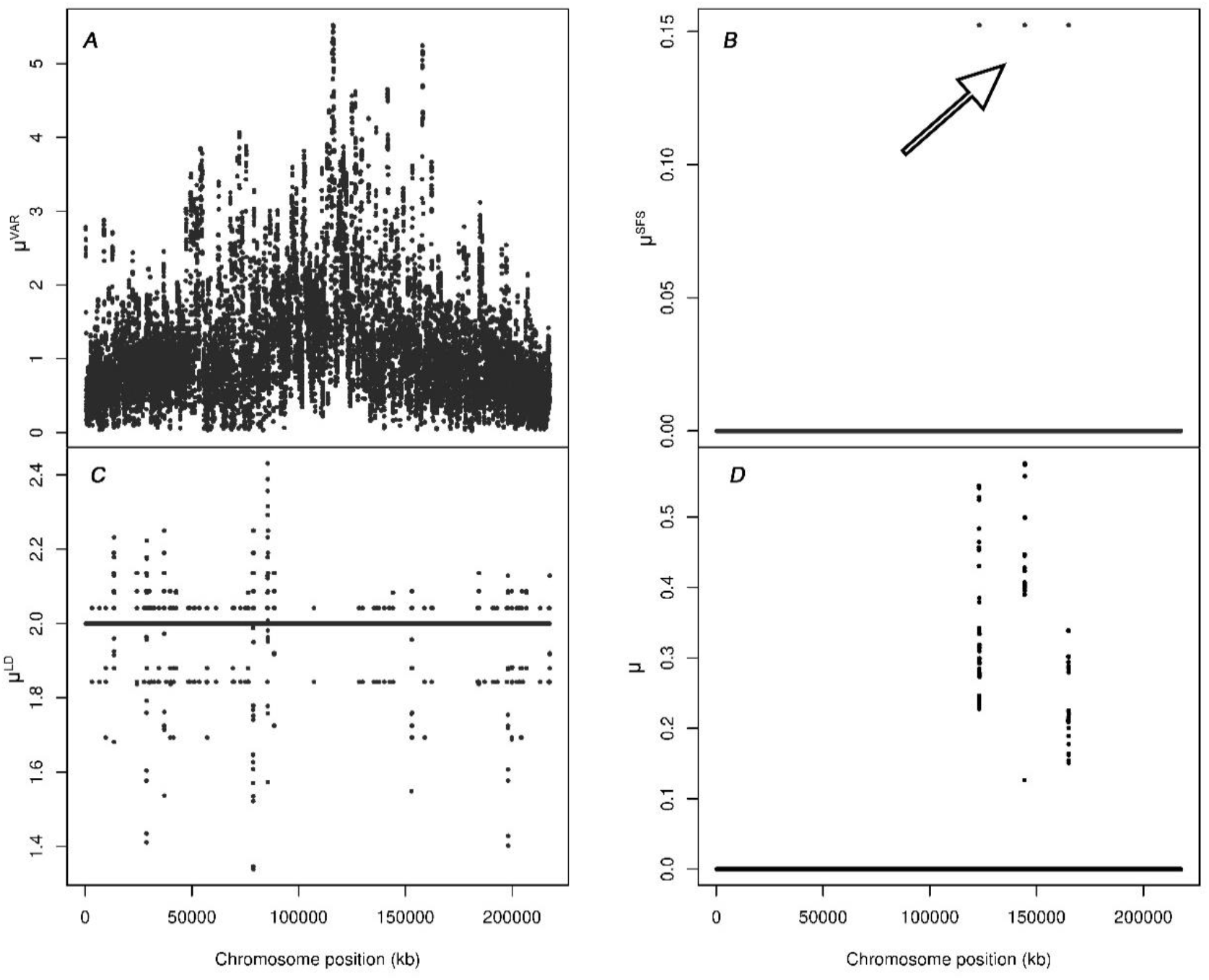
Four RAiSD *μ* parameters (Figure 6A - ◻^VAR^; Figure 6B - ◻^SFS^; Figure 6C - ◻^LD^, and Figure 6D - ◻◻◻for potential selective sweeps on chromosome 5 in the SEE maize panel.

Another selective sweep signal was detected with RAiSD on chromosome 6 (Figure 7). The values of μ^VAR^, μ^SFS^ and μ^LD^ resulted in final estimates of sweep statistics (μ) of 0.31 on start position 118933183 bp. The search for candidate genes within the regions with non-zero μ statistics was carried out within MaizeGDB interface. Within the region with start position 122994833 bp in chromosome 5, a gene coding for gras17 -GRAS transcription factor is found (Table 4). In position 144327275 bp several protein coding genes are found, namely bHLH transcription factor, putative protein phosphatase 2C 76 and Rhodanese-like domain-containing protein 4 chloroplastic. Within the last detected putative selective sweep in chromosome 5, position 164884576 bp, protein coding genes for Polyadenylate-binding protein-interacting protein 3, RS21-C6, Os02g0478550-like, rps27b and Spotted leaf protein 11 are found. In the Chromosome 6 within the region in which selective sweep signal was detected, is the protein coding gene bzip59 - bZIP-transcription factor 59, and uncharacterized genes TIDP3136 and AC209629.2_FG003.

**Figure 7.**
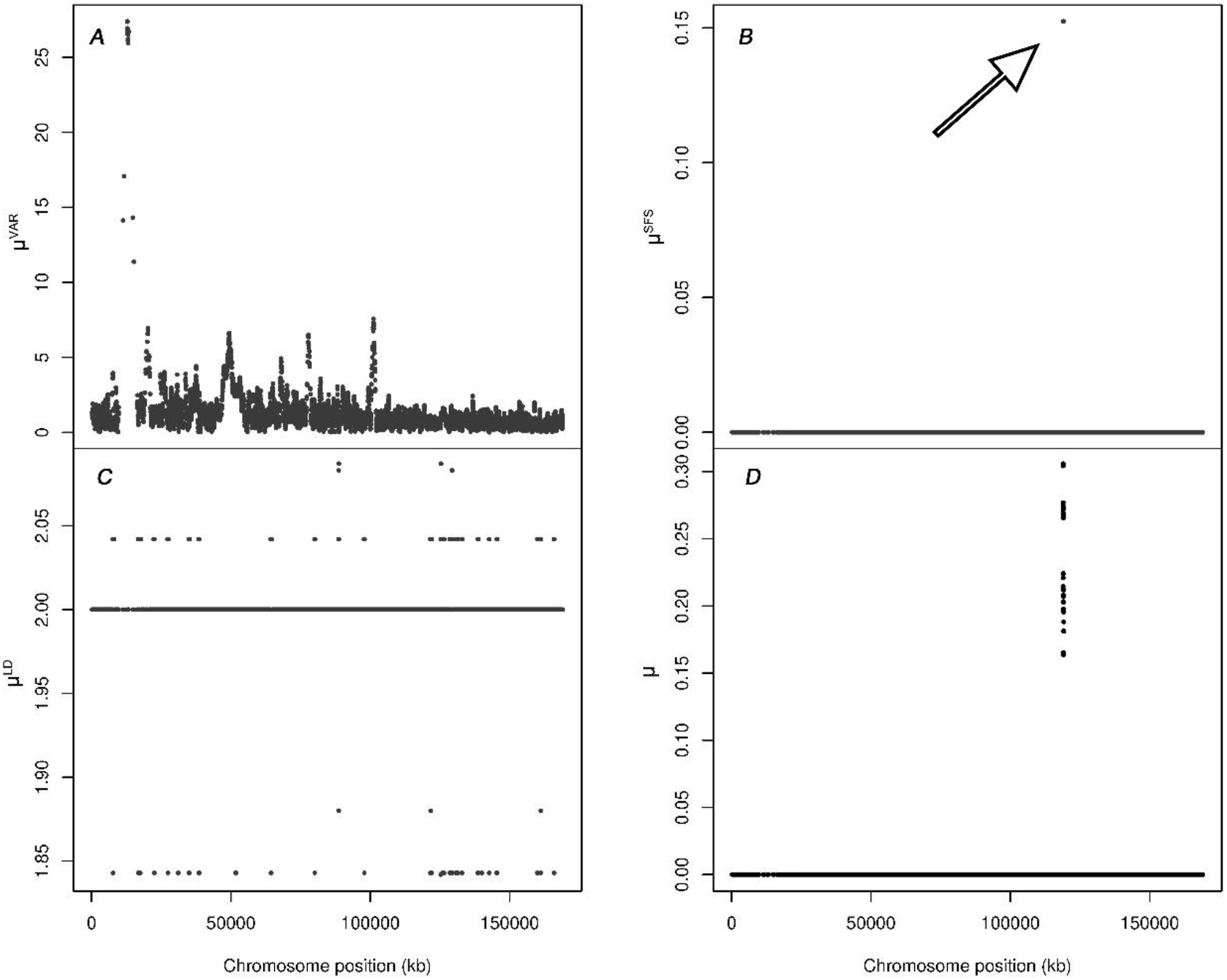
Four RAiSD *μ* parameters (Figure 7A - ◻^VAR^; Figure 7B - ◻^SFS^; Figure 7C - ◻^LD^, and Figure 7D - ◻◻◻for potential selective sweeps on chromosome 6 in the SEE maize panel.

**Table 4.** Candidate genes within potential selective sweeps on chromosomes 5 and 6

| Chromosome | Start position | Final position | $\mu$ (max) | Candidates |
| --- | --- | --- | --- | --- |
| 5 | 122994833 | 123266341 | 0.54 | gras17 -GRAS transcription factor |
| 5 | 144327275 | 144538840 | 0.58 | bHLH transcription factor 139, putative protein phosphatase 2C 76, Rhodanese-like domain-containing protein 4 chloroplastic |
| 5 | 164884576 | 165073595 | 0.34 | Polyadenylate-binding protein-interacting protein 3 (CID3), RS21-C6, Os02g0478550-like, rps27b, Spotted leaf protein 11 (SLP11) |
| 6 | 118933183 | 119093347 | 0.31 | bzip59 - bZIP-transcription factor 59, TIDP3136, AC209629.2_FG003 |

## 4. Discussion

This study represents a historical perspective on the germplasm of the SEE and provides the first information needed to successfully utilize the favorable genetic information by overcoming the issues of the classical breeding approach.

For K = 7, the joint STRUCTURE analysis of the three European panels showed one flint group and six dent groups represented notably by B73, Mo17, B14, Wf9, A374 related SEE inbreds and Iodent lines, respectively. While subpopulations K1-K4 contained inbreds belonging to all three genotyping panels, there is a clear prevalence of the lines from SEE in K5 and K6 subpopulations with admixture coefficients >0.9. The first represents the Wf9, Pa91 and Oh43 based germplasm, and the latter representing the germplasm based on “A” lines, namely A374 from Minnesota breeding programs (Schaefer and Bernardo, 2013). These two groups have already been identified earlier as separate subpopulations of the temperate maize germplasm (Hansey et al., 2011; Schaefer and Bernardo, 2013). The prevalence of these lines in SEE probably reflects the early reports on the import of the historical US germplasm after the WWII (Tavčar, 1955) and their use for breeding with locally adapted landraces (Hadi et al., 2013; Leng et al., 1962). This was also confirmed by some of the more recent studies on the genetic structure of SEE germplasm (Şuteu et al., 2013). Most of these accessions are obsolete, and are not directly present in the contemporary temperate breeding germplasm (Mikel, 2011; Romay et al., 2013) except small amounts of Wf9 and Oh43 (Coffman et al., 2020). On the other hand, the Iodent germplasm (K7) is almost completely lacking in the SEE panel. This is caused primarily by the historical nature of the SEE panel, along with the fact that the Iodent progenitor line PH207 was not publicly available until 2002 (Mikel and Dudley, 2006). It might be worthwhile to re-evaluate this resource with modern tools, especially since the local SEE landraces have been used in breeding with these accessions possibly offering certain resource of alleles for adaptation traits. This is reflected through the high allelic diversity present in this panel (Table 1) accompanied by the very high estimates of the Tajima’s *D*. High *D* values represent the effects of balancing selection (Tajima, 1989). This might have been influenced by the population contraction or possibly by the selection within the known heterotic patterns. The familiar examples of the balancing selection are heterozygote advantage (overdominance in case of heterosis) and frequency-dependent selection with rare-allele advantage. The frequency dependent selection possibly strains from the fact that the present results represent the genotyping results of a genetic resource collection in which many inbreds represent the maintained admixed accessions with local landraces.

Plotting the results on the map of Europe with kriging of dominant patterns on coordinates revealed the three different underlying patterns of the distribution of ancestry coefficients. Namely, the main pattern in the Western Europe represented by the accessions from France, Germany and Switzerland is mostly of European Flint materials which is in accordance with the results of (Bouchet et al., 2013). Another pattern was represented solely by the accessions from Italy, closely related to the Stiff Stalk Synthetic germplasm. The third pattern represented by the inbreds Wf9, Pa91 and Oh43 can be observed in the Spain, Portugal and most of Eastern and Southeastern Europe. The larger proportions of the lines associated with materials from Minnesota in SEE can also be observed (Figure 4, blue).

The scan for selective sweeps using Tajima’s *D* statistic yielded very high estimates of *D* throughout the genome. The high estimates of *D* are expected in cases of balancing selection, and heterotic patterns in maize that maximize the heterotic effects make the balancing selection inevitable, especially in commercial germplasm. However, the possible signal of a soft selective sweep was detected on chromosome 2, where a gene coding for *Flowering promoting factor-like 1* (FPF1) protein is found. FPF1 is involved in floral development and transition from vegetative to reproductive phase of plant. BLAST of the cDNA sequence gave 84-100% covers in maize, sorghum and weeping love grass (*Eragostis curvula*), possibly indicating a gene conserved in C4 grasses. Soft sweeps appear to be the signature of a main mechanism of adaptation, i.e. they do not result in a large shift in the site frequency spectrum leaving the genetic variation within position slightly changed (Luikart et al., 2018). Moreover, variation in flowering regulation provides maize the means of adaptation to different latitudes and longitudes (Bouchet et al., 2013; Romero Navarro et al., 2017) influenced by different day lengths, temperatures, and stressors (Brandenburg et al., 2017). However, another reason for this signal might be the selfing of the first pollinating progenies for many generations in breeding programs causing this putative soft sweep signal as overexpression of this gene leads to shortening of time to flowering (Wang et al., 2014). The original inbreds in complete linkage disequilibrium were probably left unaffected, thus preventing the hard sweep signal.

The further scan for selective sweeps was performed using the Raised Accuracy in Sweep Detection (RAiSD) methodology (Alachiotis and Pavlidis, 2018). RAiSD was chosen because it combines the three known signals of selective sweeps in calculation of μ statistic: local reduction of polymorphism levels, shift in the site frequency spectra, and the localized patterns of linkage disequilibrium within the 50 bp windows thus providing the increased accuracy of true positive detection of approximately 97%. The detection of a selective sweeps is under the strong influence of the migration and bottlenecks which is especially applicable to the breeding germplasm, regularly exchanged between breeders, companies and plant genetic resource offices. This can generate the large number of false positives, so defining the cutoff of at least 95% is advisable. In our work, the shown sweep signal statistics μ on chromosomes 5 and 6 (Figures 6d and 7d) both fall below the 99^th^ percentile for the individual chromosomes in which the signals were detected. It appears that all four detected sweep candidate loci were driven by the highly altered site frequency spectra (SFS, Figure 6b and Figure 7b). The changes in SFS are usually caused by the background selection for beneficial variants, which increase in frequency accompanied by the decrease in frequency of positions not linked to beneficial variants (Pavlidis and Alachiotis, 2017). All sweep signals were detected within the gene-rich regions. In the region with start position 122994833 bp in chromosome 5, GRAS transcription factor (gras17) is located. The gras17 is involved in processes of meristem initiation and regulation of transcription with highest expression levels in shoot and leave tips (Stelpflug et al., 2016). The GRAS family of transcription factors is very large with only a few characterized genes with known physiological roles (Guo et al., 2017), so it is not possible to establish the cause of background selection of one variant over other. In the second position on chromosome 5 (144327275 bp), the basic Helix-Loop-Helix (bHLH) transcription factor was detected. The most famous of bHLH transcription factor gene is a *BARREN STALK1* which regulates formation of axillary tissues including tillers (Woods et al., 2011), possibly indicating selection against tillering. The bHLH139 detected in this study is still uncharacterized, but its duplications through the genome indicate an important biological role (Zhang et al., 2018). Selective sweep might thus also indicate the selection of a single morphological type in some morphological characteristic, or adaptation to certain environmental factors. Of the candidates located in the last detected region in chromosome 5 (164884576 bp), two have overlapping roles and might have been inadvertent targets of selection. The first is CID3, coding for Polyadenylate-binding protein-interacting protein 3, involved in responses to auxin stimulus (Wada et al., 2012). The second is SPL3 (Spotted leaf protein 11), involved in flowering, with elevated expression levels in reproductive organs (Shikata et al., 2009). Although there are known roles for these two genes, some other uncharacterized gene might also have been under selection causing the detected signal. On chromosome 6, position 118933183 bp, a basic leucine zipper transcription factor 59 (bZIP59) is located with molecular function involved in DNA-binding transcription factor activity. The bZIP represents a large family of transcription factor, with some known genes included in the protein storage in grain, such as Opaque2 (Yang et al., 2016), and many factors included in the seed development which might have influenced the selection (Wang et al., 2019).

## 5. Conclusions

The distinct genetic structure patterns were detected in the SEE when genotyping results were analyzed in pan-European context provided by the two other publically available complementary European panels. Some of the prevailing ancestral patterns in historical accessions from SEE can be explained by several historical references on the import and use for breeding of certain historical inbreds, such as Wf9, Pa91, Oh43 and A374 (Tavčar, 1955). High nucleotide diversity in the SEE panel might also be partially caused by the use of local landraces in pedigrees of some inbreds (Leng et al., 1962). Soft sweep signal detected in the region of chromosome 2, harboring the gene FPF1, with known role in induction of flowering might have been caused by the extensive pollination of the first flowering progenies in crosses from which the inbreds were developed. Additional scan for selective sweeps using the RAiSD methodology yielded three more sweep signals in chromosome 5, and a single sweep signal in chromosome 6. All sweeps were detected in regions harboring genes affecting morphology and flowering, possibly indicating the inadvertent selection for the best-adapted or the favorable-appearance types. Our study provides the first step towards the utilization of this rich resource of the genetic materials for use in breeding. Accompanying phenotypic analysis is needed for assessment of the SEE accessions for favorable alleles, and identification of breeding targets.

